# Anticlastogenic Effect of *Ulva Fasciata*, Against Cyclophosphamide and Mitomycin C Induced Chromosomal Damage in Swiss Albino Mice

**DOI:** 10.1101/370031

**Authors:** Jayashree Dolpady, K.K. Vijayalaxmi

## Abstract

Chemoprevention is a strategy to reduce the incidence of human cancer either by inhibiting initiation of carcinogenesis or by preventing exposure to carcinogens, by the use of plant or animal derived ingredients. In the present study we investigated the anticlastogenic effect of ethanol extract of *Ulva fasciata*, a green seaweed, against the chromosomal aberration and micronucleus induced by the anticancer drugs cyclophosphamide and mitomycin C. Three doses of extract (25, 50 and 75 mg/kg.b.w) was given by oral gavage for 5 days at 24 hr. intervals and on the 5^th^ day, CP (25 and 50mg/kg.b.w) or MMC (1 and 2mg/kg.b.w) were intraperitoneally injected and 24hrs. later micronucleus and chromosomal aberrations assays were performed. Our results show that Ulva extract gave significant protection against the CP and MMC induced damages by reducing micronucleus and chromosomal aberrations. The protection imparted by Ulva could be due to the synergistic and/or additive effects of biologically active ingredients present in the seaweed.

## INTRODUCTION

Most cytotoxic drugs used in cancer chemotherapy are highly toxic to wide spectrum of normal cells giving rise to various side effects. Recently, much attention has been focused on chemoprevention using naturally occurring substances present in dietary and medicinal plants. Chemoprevention is a strategy used to reduce the incidence of human cancer either by inhibiting initiation and spreading of carcinogenesis or by preventing damages due to the exposure to carcinogens, by natural ingredients from plant or animal sources ^1^. Anticlastogens are the agents that can reduce the chromosomal damage induced by clastogens ^2^.

Of the abundant plant species (more than 3 lakhs) known in the world, some 40,000 species belong to Algae, including seaweeds. Seaweeds are used worldwide in food, textile, chemical, pharmaceutical and cosmetic industries. Chinese, Japanese, Pilipinos and other South East Asians use algae not only as food but also in medicine since centuries ^3^. There are plenty of edible seaweeds all along the Indian coast. Most of the seaweeds from Indian coast are exploited for the production industrial products such as agar, alginates, carrageenan and fertilizers. *Gracilaria lichenoids* is widely used in the coastal districts of Tamilnadu as porridge ^4^. Scientific research on chemopreventive effects of seaweeds are rarely been investigated. However, there are some reports on the anticlastogenic ^5^, antitumor ^6-8^, antioxidant, antimutagenic and antiproliferative ^9^ properties of seaweeds and their isolated compounds from around the world.

*Ulva fasciata,* Delile (Chlorophyceae) (henceforth referred as Ulva or extract), is commonly known as sea lettuce, grows in the intertidal rocks and abundant in the nutrient rich estuaries. In Hawai, it is a popular food eaten as salad. Chemical composition of Ulva revealed, protein, sulfate, uronic acid and sugar ^10^, xylorhamnoglucuronan and glucuronic acid ^11^, Chlorophyl, β-sitosterol, isofucosterol and saringosterol ^12^ and vitamin C ^13^.

There are a few reports available on studies such as antibacterial, antiviral, antiprotozoal and antifertility effects of Ulva ^14, 15^ and anticancer activity of Ulva ^16, 17^. These health benefits of Ulva prompted us to study the anticlastogenic properties of alcoholic extract of Ulva which was abundantly available in the Mangalore coast, that was not been studied. In this study we investigated the protective effects of Ulva against chromosomal damage induced by the well-known anticancer drugs such as cyclophosphamide (CP) and Miomycin C (MMC) in Swiss albino mice by using micronucleus and chromosomal aberration assays.

## MATERIALS AND METHODS

### Collection of seaweeds and extraction

Ulva was collected during the months of August to November (2001-2002) from Someshwara beach, Dakshina Kannada District, Karnataka. The seaweed was identified using identification keys18 and confirmed by department of Applied Botany Mangalore University (Voucher no. SW02).The collected seaweed was freed from extraneous debris, sand and other associated macro-fauna, washed thoroughly and shade dried. Dried seaweed was ground into powder with a mixer. 100gm of powdered seaweeds was extracted with 500ml of distilled ethanol in a Soxhlet extractor for about 8hrs. The extract thus obtained was concentrated in a rotary vacuum evaporator (Laborota 4003, Heidolph Rotavac, Germany) at 40°C and thick paste was obtained by evaporating in a water bath at 45°C. The yield was 1.8% and the extract was kept in the deep freezer for further use.

### Toxicity study

The extract was insoluble in water hence used 10% alcohol as solvent. The toxicity of extract was tested in Swiss albino mice by following the method by Ghosh ^19^. Mice were divided into different groups of 10 animals (5 males & 5 females in each group). Various doses of the extract, viz. 100, 200, 300, 400, 500, 1000, 2000, 3000, 4000 and 5000 mg/kg body weight were prepared in 10% alcohol and administered to different groups of fasting animals through oral gavage method. After administering the extract, feeding was resumed and animals were kept under observation for 72 hours.

### Experimental animals

Swiss albino mice (*Mus musculus*) were used for the study. The animals were housed in the shoe box cages bedded with rice husk, bred and maintained at temperature controlled animal facility of Applied Zoology department, Mangalore University. Animals were fed commercial pelleted food and water *ad libitum*. All the experimental protocols and animal handling procedures were approved by the Institutional Animal Ethical Committee (IAEC) of Mangalore University and followed as per the guidelines of CPCSEA India (232/CPCSEA Dt:01.08.2000). For each experimental groups five animals (of either sex, 25±2gms, 10-12 weeks old) were used.

### Treatment procedure

The selected doses, 25, 50 and 75mg/kg.b.w. of extract in 0.2 ml, administered by oral gavage to experimental animals for 5 days with an interval of 24 hours. The positive mutagen CP (CAS No: 6055-19-2, Batch No: 182, Endoxan, German remedies Ltd, Goa, India) or MMC (CAS No: 50-07-7, Batch No: LB 723002, Biochem. Pharmaceutical Industries, Bombay, India) was administered on the 5th day, I hour after the last extract treatment. Bone marrow micronucleus slides and metaphase chromosome slides were prepared by following the standard protocols. Animals administered with 0.2 ml of drinking water and 0.2 ml of 10% alcohol formed the control and solvent controls respectively. Positive mutagens, CP (25 and 50mg/kg, b.w.) and MMC (1 and 2 mg/ kg.b.w.) were dissolved in sterile distilled water and administered intraperitoneally to the experimental animals as single dose in 0.1 ml. All the chemicals used in the experiments were purchased from either Sigma, Merck, SRL, Himedia, India. Table 1 shows the treatment groups.

**TABLE 1:** TREATMENT, DOSE AND GROUPS USED FOR MICRONUCLEUS ASSAY AND CHROMOSOMAL ABERRATION ASSAY.

| **Treatment** | **Dose** | **Group** |
| --- | --- | --- |
| **Water** | 0.2ml-5days | Control |
| **10% alcohol** | 0.2ml-5days | Solvent control |
| **Extract** | 25,50,75 (5days) | Extract |
| **CP** | 25, 50 (single dose) | Positive mutagen |
| **Extract + CP** | E25+CP25, E25+CP50 | Combination treatment |
|  | E50+CP25, E50+CP50 | Combination treatment |
|  | E75+CP25, E75+CP50 | Combination treatment |
| **MMC** | 1,2 (single dose) | Positive mutagen |
| **Extract+ MMC** | E25+M1, E25+M2 | Combination treatment |
|  | E50+M1, E50+M2 | Combination treatment |
|  | E75+M1, E75+M2 | Combination treatment |

### Micronucleus (MN) assay

Bone marrow MN slides were prepared by following the modified method of Schmid ^20^ by using bovine serum albumin ^21^. Bone marrow cell suspension in 3% BSA was centrifuged at 1000RPM. The pellet was suspended in suspension medium, a drop of the suspension was smeared on a clean slide and air-dried, fixed in methanol and stained with May-Grunwalds-Giemsa. 2000 Polychromatic Erythrocytes (PCE) and Normochromatic Erythrocytes (NCE) in the corresponding field were screened for each animal to score MN and to determine the ratio of P/N.

### Chromosome aberration (CA)Assay

The preparation of bone marrow metaphase chromosome was prepared by following the method of Tjio & Whang ^22^. Experimental mice were injected i.p. with 0.024% colchicine (0.2ml) after the drug treatment. Bone marrow cells were collected after 1 ½ hours later and suspended in 0.56%KCl and kept for 20 minutes, centrifuged, re-suspended the pellet in 2ml fixative (1:3 acetic acid-methanol for 45 minutes) and centrifuged again. Then the suspension was dropped onto pre-chilled slides, flame dried and stained with Giemsa. For scoring aberration, the slides were screened under 1000X (Olympus BX). One hundred well spread metaphases were examined per each animal and total of 2000 dividing and non-dividing cells for each animal was also screened on the same spots in order to determine the mitotic index. Aberrations like break, rings, centric fusion, exchange, fragmentation and multiple aberrations were taken into consideration.

The results were expressed as mean ± standard error. Differences between the groups were analyzed by paired T test ^23^.

## RESULTS

### Toxicity study

We observed no toxicity signs in the tested doses in the initial 72 hrs. The animals were further observed for 14 days to determine the toxic/lethal effects of the extract. None of these doses induced mortality or any behavioral changes such as grooming, tremors, convulsion etc. observed in the experimental animals. These results indicated the tested doses were safe hence we selected doses 25, 50 and 75 mg/kg. b.w. for the study.

Both the doses of CP and MMC induced significant (p< 0.001) frequency of micronuclei in polychromatic erythrocytes (PCE) and total MN compared to control groups. P/N ratio was reduced in CP and MMC treated animals. CP and MMC also induced significant (p< 0.001) amount of chromosomal aberrations in both the doses tested compared to the control groups.

### Effect of Ulva extract on Micronucleus

Ulva extract alone did not induce significant MN at any of the doses tested. With CP, Ulva extract imparted significant (p< 0.001) protective effects by reducing the MN in all the three doses in a dose dependent manner. With MMC also the protective effect of the extract was significant, although the percent inhibition was not as much as that observed with CP. Ulva extract also enhanced the P/N ratio in the CP treated animals which was found to be significant (p< 0.001 and p< 0.01) compared to MMC at certain instances (p< 0.05). Results are presented in Table 2.

**TABLE 2:** EFFECT OF ETHANOLIC EXTRACT OF ULVA FASCIATA ON MN INDUCED BY CP AND MMC IN MOUSE BONE MARROW CELLS.

| **Treatment (mg/kg.b.w)** | **MNPCE<sup>A</sup> ±SEM(%)** | **Inhibition (%)** | **Total MN ±SEM (%)** | **Inhibition (%)** | **P/N ± SEM (%)** |
| --- | --- | --- | --- | --- | --- |
| **Water** | 0.18±0.23 | - | 0.14±0.01 | - | 1.01±0.00 |
| **10%Alc** | 0.23±0.02 | - | 0.11±0.01 | - | 1.02±0.02 |
| **E 25** | 0.30± 0.02 | - | 0.18±0.02 | - | 0.97±0.01 |
| **E 50** | 0.32±0.03 | - | 0.22±0.04 | - | 1.10±0.02 |
| **E 75** | 0.25±0.06 | - | 0.15±0.02 | - | 1.05±0.04 |
| **CP 25** | 1.21±0.06 <sup>c</sup> |  | 0.64±0.03 <sup>c</sup> |  | 0.73±0.05 <sup>c</sup> |
| **E25+CP25** | 0.92±0.06 <sup>a</sup> | 29.59 | 0.50±0.03 <sup>a</sup> | 26.42 | 0.92±0.02 <sup>b</sup> |
| **E50+CP25** | 0.67±0.05 <sup>c</sup> | 52.92 | 0.41±0.02 <sup>b</sup> | 43.39 | 1.13±0.01 <sup>c</sup> |
| **E75+CP25** | 0.49±0.11 <sup>c</sup> | 73.47 | 0.29±0.07<sup>c</sup> | 66.04 | 1.09±0.02 <sup>c</sup> |
| **CP 50** | 2.16±0.05 <sup>c</sup> |  | 1.19±0.05 <sup>c</sup> |  | 0.59±0.01 <sup>c</sup> |
| **E25+CP50** | 1.18±0.10 <sup>c</sup> | 50.78 | 0.58±0.10 <sup>c</sup> | 56.48 | 0.76±0.16 <sup>b</sup> |
| **E50+CP50** | 0.65±0.08 <sup>c</sup> | 78.24 | 0.36±0.03 <sup>c</sup> | 76.85 | 0.91±0.02 <sup>c</sup> |
| **E75+CP50** | 0.27±0.07 <sup>c</sup> | 97.93 | 0.15±0.03 <sup>c</sup> | 96.29 | 0.81±0.01 <sup>c</sup> |
| **MMC 1** | 1.04±0.05 <sup>c</sup> |  | 0.51±0.04 <sup>c</sup> |  | 0.77±0.01 <sup>c</sup> |
| **E25+MMC1** | 0.81±0.06 <sup>a</sup> | 28.39 | 0.40±0.02 <sup>a</sup> | 27.50 | 0.80±0.02 |
| **E50+MMC1** | 0.72±0.06 <sup>a</sup> | 39.51 | 0.37±0.02 <sup>a</sup> | 35.00 | 0.94±0.17 <sup>a</sup> |
| **E75+MMC1** | 0.62±0.15 <sup>c</sup> | 51.85 | 0.32±0.07 <sup>b</sup> | 47.50 | 0.95±0.02 <sup>a</sup> |
| **MMC 2** | 2.00±0.13 <sup>c</sup> |  | 1.01±0.04 <sup>c</sup> |  | 0.79±0.01 <sup>c</sup> |
| **E25+MMC2** | 1.63±0.21 | 20.90 | 0.78±0.07 <sup>a</sup> | 25.56 | 0.80±0.01 |
| **E50+MMC2** | 1.50±0.12 <sup>a</sup> | 28.25 | 0.72±0.05 <sup>a</sup> | 32.20 | 0.82±0.01 |
| **E75+MMC2** | 1.23±0.10 <sup>b</sup> | 43.50 | 0.64±0.06 <sup>b</sup> | 41.11 | 0.95±0.03 <sup>a</sup> |

<sup>A</sup> - 2000 PCE from each animal (5 animals / group), 't' test: <sup>a</sup> p< 0.05, <sup>b</sup> p<0.01, <sup>c</sup> p<0.001
PCE=Polychromatic Erythrocytes, MN=Micronucleus
P/N = Polychromatic Erythrocytes Normochromatic Erythrocytes

### Effect of Ulva extract on chromosomal aberrations

Total aberrations with two doses of CP were 48.8 % and 55.6 % and with two doses of MMC were 29.7% and 42.7% respectively which was dose dependent. Total percent aberrations induced by 10% alcohol was insignificant. Ulva extract alone did not induce any significant chromosomal aberrations in the selected three doses. In the combination treatment with the positive mutagens, Ulva extract imparted protective effects, as manifested by the reduction in the chromosomal aberrations. The protective effect was found to be more significant (p< 0.001) in the combination of higher dose of extract +lower dose of drug. The results are presented in Table 3. Extract 50 and 75 mg doses with CP 25mg showed more than 60% inhibition in the chromosomal aberration. Extract 75mg dose gave better protective effect with CP 50mg, MMC 1 and 2mg doses. The results are presented in the Figures 1 and 2.

**TABLE 3:** EFFECT OF ETHANOLIC EXTRACT OF ULVA FASCIATA ON CA INDUCED BY CP AND MMC IN MOUSE BONE MARROW CELLS.

| **Treatment**<br>**(mg/kg.b.w)** | **MI** | **NC** | **BS** | **R** | **CF** | **EX** | **F** | **MA** | **TA** |
| --- | --- | --- | --- | --- | --- | --- | --- | --- | --- |
| **Water** | 5.1±0.02 | 98.50 | 0.4 | 0.5 | 0.6 | - | - | - | 1.5±0.28 |
| **10%Alc** | 5.1±0.08 | 94.70 | 2.0 | 1.9 | 1.2 | 0.2 | - | - | 5.3±3.20 |
| **E 25** | 4.9±0.13 | 97.50 | 1.9 | - | 0.6 | - | - | - | 2.5±0.31 |
| **E 50** | 4.9±0.02 | 96.10 | 2.3 | 0.4 | 1.2 | - | - | - | 3.9±2.10 |
| **E 75** | 4.8±0.09 | 97.50 | 2.3 | - | 0.2 | - | - | - | 2.5±3.10 |
| **CP 25** | 4.5±0.13 | 52.21 | 21.3 | 4.1 | 3.1 | 10.0 | 2.7 | 7.6 | 48.8±1.16<sup>c</sup> |
| **E25+CP25** | 4.4±0.04 | 67.91 | 15.4 | 2.6 | 1.6 | 5.9 | 0.8 | 5.8 | 32.1±0.71<sup>c</sup> |
| **E50+CP25** | 4.6±0.14 | 77.40 | 15.8 | 1.7 | 1.1 | 1.5 | 0.6 | 1.9 | 22.6±1.89<sup>c</sup> |
| **E75+CP25** | 4.8±0.09<sup>a</sup> | 76.84 | 13.8 | 1.6 | 1.2 | 3.6 | 0.6 | 2.4 | 23.3±0.96<sup>c</sup> |
| **CP 50** | 3.6±0.17<sup>c</sup> | 44.40 | 25.1 | 2.3 | 2.1 | 6.7 | 12.1 | 7.3 | 55.6±0.76<sup>c</sup> |
| **E25+CP50** | 5.1±0.16<sup>c</sup> | 51.32 | 21.4 | 2.7 | 2.1 | 5.2 | 10.5 | 6.8 | 48.7±0.66<sup>b</sup> |
| **E50+CP50** | 4.8±0.08<sup>c</sup> | 55.33 | 20.7 | 2.2 | 2.0 | 5.7 | 8.8 | 5.3 | 44.7±1.02<sup>c</sup> |
| **E75+CP50** | 5.1±0.12<sup>c</sup> | 60.00 | 19.4 | 2.4 | 2.0 | 5.3 | 6.3 | 4.6 | 40.0±0.79<sup>c</sup> |
| **MMC 1** | 4.7±0.06<sup>c</sup> | 70.30 | 18.8 | 2.5 | 1.9 | 4.6 | - | 1.9 | 29.7±0.87<sup>c</sup> |
| **E25+MMC1** | 5.5±0.32<sup>a</sup> | 78.12 | 18.6 | 1.9 | 2.7 | 4.4 | 0.2 | 1.1 | 28.9±0.22 |
| **E50+MMC1** | 5.4±0.09<sup>a</sup> | 73.74 | 17.2 | 2.5 | 2.9 | 2.9 | - | 0.8 | 26.3±1.38<sup>b</sup> |
| **E75+MMC1** | 5.5±0.06<sup>a</sup> | 78.60 | 15.1 | 2.0 | 1.9 | 1.5 | 0.2 | 0.7 | 21.4±0.93<sup>c</sup> |
| **MMC 2** | 3.9±0.06<sup>c</sup> | 57.30 | 27.0 | 2.1 | 2.3 | 4.1 | 1.8 | 5.4 | 42.7±0.81<sup>c</sup> |
| **E25+MMC2** | 4.2±0.08 | 62.40 | 19.7 | 2.3 | 2.7 | 7.0 | 2.0 | 3.9 | 37.6±0.96<sup>b</sup> |
| **E50+MMC2** | 4.4±0.06<sup>a</sup> | 68.30 | 19.7 | 2.7 | 1.6 | 3.2 | 1.6 | 2.9 | 31.7±1.92<sup>c</sup> |
| **E75+MMC2** | 4.6±0.06<sup>b</sup> | 73.90 | 16.6 | 2.8 | 1.6 | 3.2 | 0.4 | 1.5 | 26.1±0.88<sup>c</sup> |

A=2000 cells/animal (5 animals/group)
MI= Mitotic Index, BS=Break, R=Ring, CF=Centric fusion,
EX=Exchange, F=Fragmentation, MA=Multiple aberration
‘t’ test: <sup>a</sup> p< 0.05, <sup>b</sup> p<0.01, <sup>c</sup> p<0.001

**FIGURE 1:**
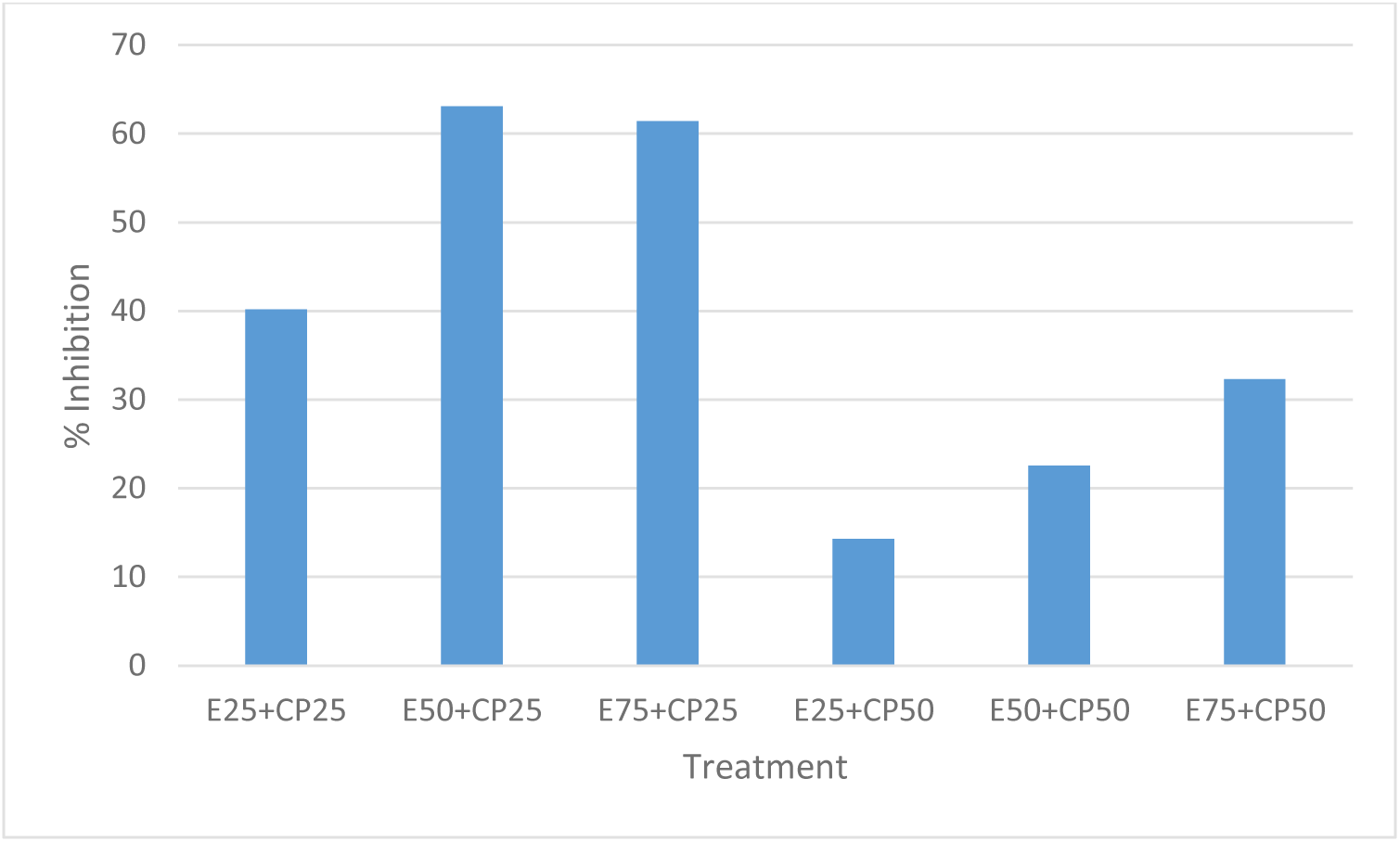
PERCENT INHIBITION OF TOTAL CHROMOSOMAL ABERRATIONS BY ULVA EXTRACT IN CP TREATED GROUPS.

**FIGURE 2:**
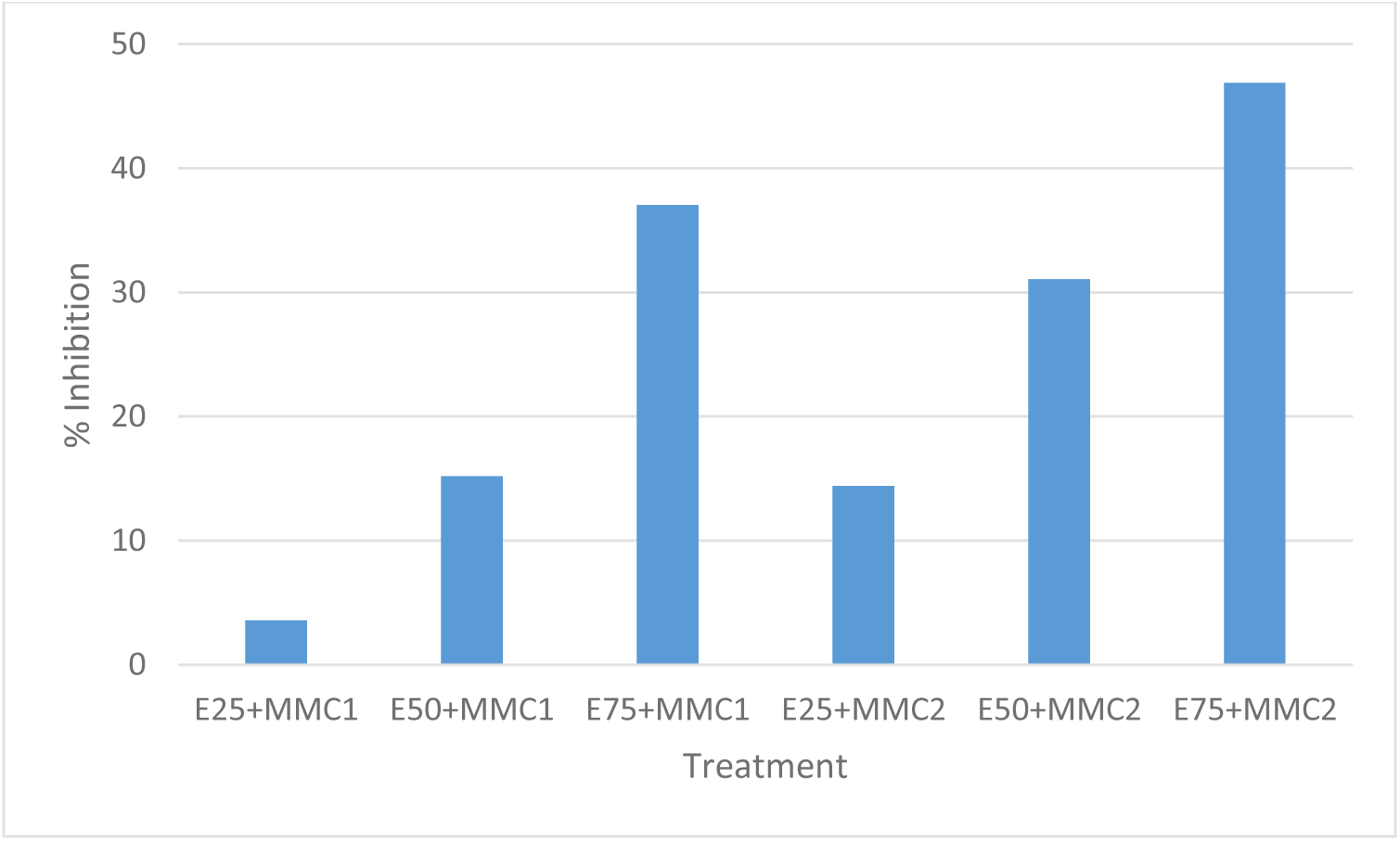
PERCENT INHIBITION OF TOTAL CHROMOSOMAL ABERRATION BY ULVA EXTRACT IN MMC TREATED GROUPS.

## DISCUSSION AND CONCLUSION

The ideal chemotherapeutic drug for cancer should destroy only the cancer cells without any adverse effects or toxicities to normal cells. But, so far no such drugs have been identified. CP and MMC which are used as standard anticancer drugs for various types of cancer are also used as standard drug to induce micronucleus and chromosomal aberrations in the anticancer research experiments.

In the present study, Ulva extract showed protective effects against the clastogenicity of CP and MMC which was evaluated by micronucleus test and chromosomal aberration assay. Cytotoxicity of CP and MMC was evidenced in the present study in the form of reduction in the mitotic index (Ml) and P/N ratio. Ulva extract at different doses enhanced the P/N ratio in MN study and Ml in CA study at certain instances which may be due to the stimulatory action of this extract on the cell division. In micronuclei test the combination treatment of extract with CP or MMC showed protection in a dose dependent manner. Maximum protection was observed at CP 50mg and extract 75mg. In case of chromosomal aberration also we could observe similar results that higher doses of extract significantly reduced total chromosomal aberration induced by CP and MMC. In both MN and CA test we observed CP was more effective in imparting protection compared to MMC. This can be explained that both CP and MMC belong to different class of drugs and their mechanism of interaction is also different. Our findings are in correlation with the earlier reports in which researchers observed in vivo protective effects of *Spirulina fusiformis* against CP and MMC induced genotoxicity and reported significant reduction in the chromosomal aberrations after treatment with algal extract ^24^. In another study, Blue green alga *Chloreila vulgaris* reduced X-ray induced chromosomal damage in whole body irradiated mice ^25^. Recent study on in vitro antigenotoxicity of *Ulva rigida* extract showed protection against induction of chromosome aberration, sister chromatid exchange and micronuclei by mutagenic agent MMC ^26^.

We presume that the protective effects imparted by Ulva may be due to the synergistic action of phytochemicals present in the extract. The different constituents of natural products interact additively or synergistically leading to significant antigenotoxic effects 27-29. As mentioned above, Ulva consists of phytochemicals such as chlorophyll, vitamins, flavonoids, beta-sitosterol etc. We understand that the nutrients and chemical composition may vary with the seasonal changes and geographical locations. These constituents individually have been tested for their antimutagenic and anticancer properties. Anticlastogenecity of chlorophyllin in different cell cycle phases in cultured mammalian cells was studied using chromosomal aberrations as the experimental end point. It reduced the DNA damage induced by ethyl methane sulphonate (EMS) and the effect was cell cycle dependent ^30^. Chlorophyll a and b showed protection against MMC induced chromatid and isochromatid breaks in human lymphocyte cultures ^31^. Sarkar et al ^32^ reviewed the chlorophyll and chlorophyllin as modifiers of genotoxic effects.

There are several reports on the antimutagenic and anticlastogenic properties of vitamins especially the vitamin-C, A and E. Vitamin-C mediated protection on cisplatin induced mutagenicity has been reported using bone marrow micronucleus and chromosomal aberrations assays ^33^. Vitamin-C, E and beta-carotene showed reduction in DNA damage induced by gamma-ray in human lymphocytes ^34,35^. In another study it was reported that vitamin-C and E induced reduction in rifampicin induced chromosomal aberrations in mouse bone marrow cells ^36^. Beta sitosterol is gaining importance these days on account of its wide range of biological activity. Villasenor et al.^37^ reported that beta-sitosterol from *Mentha cordifolia* inhibited the mutagenicity of tetracycline in an in vivo MN study. In another study beta sitosterol reduced significantly the sister chromatid exchanges and MN induced by doxorubicin, it also showed reduction in the free radicals thereby giving cellular protection ^38^. These active ingredients might act at cellular level by enhancing the enzymatic action in order to detoxify the mutagens or inhibit the enzymes which take part in the formation of metabolites or scavenging the free radicals ^39^. The exact mechanism of protection is not understood; we assume that extract may or may not acting directly on the drugs or on its metabolites instead the phytochemicals present in the extract provides cyto-protective environment to the cells in order to protect from the damages caused by these drugs. Hence, we claim that using Ulva in diet may bring added benefits to the patients undergoing treatments with anticancer drugs such as CP and MMC. We conclude that further research on antitumor or antioxidant effects of Ulva extract needed to understand it’s health benefits which will lead to explore individual active ingredient with potential human therapeutic value.

## ACKNOWLEDGEMENT

The author wish to express sincere thanks to the supervisor Late Professor K.K. Vijayalaxmi. This work was carried under the project sanctioned to KKV by Department of Biotechnology, Government of India. JD wish to thank DBT for financial support to carry out this research.

